# Kilometer-scale larval dispersal processes predict metapopulation connectivity pathways for *Paramuricea biscaya* in the northern Gulf of Mexico

**DOI:** 10.1101/2021.10.06.463363

**Authors:** Guangpeng Liu, Annalisa Bracco, Andrea M. Quattrini, Santiago Herrera

## Abstract

Fine-scale larval dispersal and connectivity processes are key to species survival, growth, recovery and adaptation under rapidly changing disturbances. Quantifying both are required to develop any effective management strategy. In the present work, we examine the dispersal pattern and potential connectivity of a common deep-water coral, *Paramuricea biscaya*, found in the northern Gulf of Mexico by evaluating predictions of physical models with estimates of genetic connectivity. While genetic approaches provide estimates of realized connectivity, they do not provide information on the dispersal process. Physical circulation models can now achieve kilometer-scale resolution sufficient to provide detailed insight into the pathways and scales of larval dispersal. A high-resolution regional ocean circulation model is integrated for 2015 and its advective pathways are compared with the outcome of the genetic connectivity estimates of corals collected at six locations over the continental slope at depths comprised between 1000 and 3000 meters. Furthermore, the likely interannual variability is extrapolated using ocean hindcasts available for this basin. The general connectivity pattern exhibits a dispersal trend from east to west following 1000 to 2000-meter isobaths, corresponding to the overall westward near-bottom circulation. The connectivity networks predicted by our model were mostly congruent with the estimated genetic connectivity patterns. Our results show that although dispersal distances of 100 km or less are common, depth differences between tens to a few hundred meters can effectively limit larval dispersal. A probabilistic graphic model suggests that stepping-stone dispersal mediated by intermediate sites provides a likely mechanism for long-distance connectivity between the populations separated by distances of 300 km or greater, such as those found in the DeSoto and Keathley canyons.

## Introduction

Deep-water or cold-water corals are long-lived and slow-growing organisms commonly found at depths greater than 50 m (Cairns 2007; Roark et al., 2009; Sherwood and Edinger, 2009). They play an essential role in providing habitats for a diversity of vertebrate and invertebrate species and are highly susceptible to natural and anthropogenic disturbances (Guinotte et al., 2006; White et al. 2012; Hoegh-Guldberg et al., 2017; Turley et al., 2007). Understanding larval dispersal and connectivity patterns of deep-water corals is a first, necessary step for their conservation and management in response to the multiple threats they face (Botsford et al., 2009; Cowen et al., 2007; Palumbi, 2003).

While shallow corals are generally well sampled, direct surveys of deep corals are scarce because of the substantial cost and logistical difficulties (Doughty et al., 2014; Girard et al., 2019; Quattrini et al., 2015). The integration of biological data and physical ocean models has helped predict coral habitat suitability (Hu et al., 2020; Kinlan et al., 2020; Tong et al., 2013) and identify larval dispersal and connectivity patterns (Bracco et al., 2019; Breusing et al., 2016; Cardona et al., 2016; Etter and Bower, 2015; Fobert et al., 2019; Gary et al., 2020; Hilario et al., 2015; Nolasco et al., 2018; Ross et al., 2020; Storlazzi et al., 2017). Recent studies suggest that this biophysical framework offers meaningful predictions of connectivity (Gary et al., 2020; Ross et al., 2020), despite the uncertainties related to the sparsity of in-situ measurements and model biases in the representation of bottom boundary layer dynamics (see Bracco et al., 2020 for a recent review pertinent to the Gulf of Mexico).

Numerous shallow and deep-water corals populate the northern Gulf of Mexico (GoM) and contribute to the functionality and biodiversity of marine ecosystems (Cordes et al., 2008; Gil-Agudelo et al., 2020; Precht et al., 2014). Deep-water corals in the GoM are subject to various natural and anthropogenic stresses such as increasing water temperatures, acidification, overfishing, and pollution. For example, the 2010 Deep-water Horizon (DWH) oil spill released ~ 4.1 million barrels of oil into the Gulf (McNutt et al., 2012) and dramatically impacted the vulnerable coral communities in the proximity of the spill site (Fisher et al., 2014; Girard et al., 2019; White et al. 2012). This event and its ecological consequences pointed to the need for restoration actions and improved management strategies, raising interest for a better understanding of the biological and physical processes that affect coral connectivity in the deep-sea.

For deep-water corals, both mesoscale eddies (10 to 200 Km) and submesoscale circulations (1 to 10 Km), together with bottom boundary layer turbulence, influence their dispersal (Bracco et al., 2019; Cardona et al. 2016). Near-bottom submesoscale circulations such as fronts, vorticity filaments and small eddies, form due to instabilities induced by shear layers. These circulations can isolate larvae by trapping them inside their cores and transport them to other locations along the continental slope (Bracco et al., 2016). Additionally, the large vertical velocities and diapycnal mixing associated with submesoscale motions can contribute to the vertical transport of the larvae (Bracco et al., 2018; Vic et al., 2018).

Despite the growing number of studies focusing on biophysical dispersal models, connectivity studies that combine genetic data with models resolving the physical circulation and bathymetry at kilometer-scale resolution (submesoscale) are scarce (Bracco et al., 2019; Cardona et al., 2016; Fobert et al., 2019; Gary et al., 2020; Nolasco et al., 2018; Ross et al., 2020). This scarcity is due to the high computational costs of high-spatial resolution models, which limit them short temporal scales (days to months), and the challenges in obtaining sufficiently large sample sizes of deep-sea species for population genetics.

In this work we focus on *Paramuricea biscaya*, an octocoral in the family Plexauridae. *Paramuricea biscaya* is one of the most common and abundant corals in the GoM between 1200 and 2500 m (Doughty et al., 2014). Populations of this species were directly impacted by the 2010 Deep-water Horizon oil spill (DWH), particularly in the Mississippi Canyon area (White et al. 2012; Fisher et al. 2014), and thus are considered primary targets for restoration (Deep-water Horizon Natural Resource Damage Assessment Trustees, 2016). We investigate the metapopulation connectivity of *P. biscaya* in the northern GoM using a submesoscale permitting ocean circulation and larval dispersal model. We also evaluate the performance of the model by comparing potential connectivity probabilities with genetic connectivity estimates.

This paper is a companion to the paper by Galaska et al. (submitted) that describes the analyses of genetic connectivity and seascape genomics. Here, we compare the modeled current velocities to those of mesoscale resolving HYCOM-NCODA reanalysis. Furthermore, we explore the factors controlling the larval dispersal pathways and connectivity networks at the sites where *P. biscaya* occurs, in off-line Lagrangian particle integrations and on-line Eulerian dye simulations through spatial density analysis and a probabilistic graphic model. We also evaluate the potential role of intermediate populations predicted by habitat suitability models (Georgian et al., 2020) as stepping-stones for dispersal. Finally, we discuss the role of annual and inter-annual seasonality in modulating *P. biscaya* connectivity patterns in the GoM by means of a coarser mesoscale resolving data-assimilative hindcast.

## Data and Methods

### Large-scale circulation of the study area

Our study area comprises the region of the northern Gulf of Mexico that includes six sites, named after the lease blocks, where *Paramuricea biscaya* populations are known. These sites are: DeSoto Canyon 673 (DC673), Mississippi Canyon 344 (MC344), Mississippi Canyon 297 (MC297), Mississippi Canyon 294 (MC294), Green Canyon 852 (GC852), and Keathley Canyon 405 (KC405) (Doughty et al., 2014; Girard et al., 2019; Vohsen et al., 2020) (Figure 1). The 2010 Deepwater Horizon oil spill directly impacted *P. biscaya* populations at the Mississippi Canyon sites MC294, MC297, and MC344 (Fisher et al., 2014; White et al., 2012).

**Figure 1.**
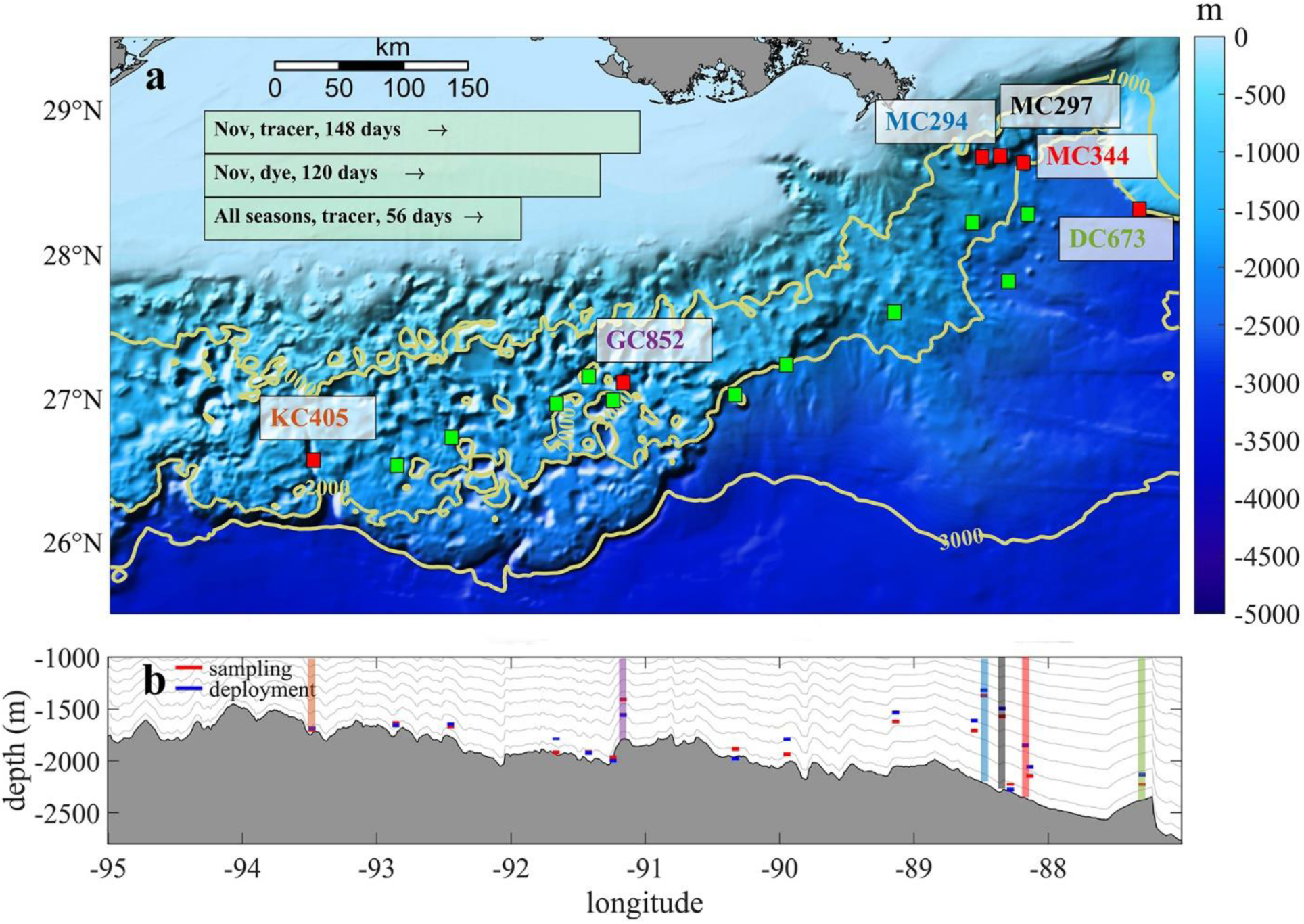
(a) Topography map of the study area showing the sampled and predicted sites hosting *P. biscaya* populations. (a) Red boxes (0.05 × 0.05º) with colored textbox indicates six main sites (from east to west, DC673, MC344, MC297, MC294, GC852, and KC405). Green boxes with black edge are predicted intermediate suitable habitats selected from models by Georgian et al. (2020). Yellow contours indicate 1000m, 2000m, and 3000m isobaths. Light green boxes at the up left corner summary three release strategies in this work. (b) The grey shaded area shows the topographic profile of the northern GoM averaged between 1000 m – 3000 m. The colored boxes indicate the depth where corals were collected (red) or larval particles released (blue). Grey lines show the average depth of the s-layers in our model.

The large-scale circulation of the study area (shown in Figure 1) is dominated in the upper 800 – 1000 m by the presence of the anticyclonic Loop Current (LC) that enters the basin through the Yucatan Channel and leaves through the Florida Straits. The LC penetrates northward to about 26.5–27.5°N and usually extends longitudinally east of 86°W (Vukovich, 1988; 2007). Large anticyclonic mesoscale eddies with diameters of about 200 kilometers spin off the main LC at irregular intervals and populate the basin until they dissipate by interacting with the continental slope in the western GoM (Cardona and Bracco, 2016; Donohue et al., 2016). At depth, below 1000 m, the large-scale circulation is cyclonic (DeHaan and Sturges, 2005; Weatherly et al., 2005). Along the continental slope, bottom currents are highly variable, and can intensify due to vortex stretching and topographic Rossby waves (Hamilton, 2009; Kolodziejczyk et al., 2012; Bracco et al., 2016).

### Larval dispersal model

Predicting larval dispersal is complicated by the limited knowledge of larvae’s behaviors, especially for deep-sea corals, and by strong, often poorly characterized, variability in deep ocean currents. The application of an integrated biophysical model remains a practicable approach to address this challenge, even though there are notable differences in the estimation of larval travel distance and dispersal pattern among different models (Cowen et al., 2007; Edmunds et al., 2018; Ross et al., 2020; Werner et al., 2007). A typical modeling framework for connectivity studies includes an ocean physical model that provides circulation, or sometimes temperature and salinity, information as background forcing field, and a module for particle (i.e., larvae) tracking those accounts for behavioral characteristics (e.g., age, life span, swimming abilities, larval buoyancy).

In this work, we adopted the three-dimensional Coastal and Regional Ocean Community model (CROCO) that is built upon the Adaptive Grid Refinement in Fortran (AGRIF) version of the Regional Ocean Modeling System (ROMS) (Debreu et al., 2012; Shchepetkin and McWilliams, 2005). It is a split-explicit, hydrostatic, and terrain-following model that is designed for simulating high-resolution nearshore and offshore dynamics and has been used successfully in larval dispersal studies (Bani et al., 2020; Bracco et al., 2019; Cardona et al., 2016; Kim and Barth, 2011; Nolasco et al., 2018; Vic et al., 2018). Here, CROCO covers a large portion of the GoM between 98°-82° W and 24°-31° N, and has a grid resolution of about 1 km in the horizontal space and 50 sigma layers in the vertical direction. The nonlinear K-Profile Parameterization (KPP) scheme parameterizes vertical mixing (Large et al., 1994). Three-dimensional tracer advection is achieved through a split and rotated 3rd-order upstream-biased advection scheme, which minimizes spurious diapycnal mixing but does not guarantee positive values of tracer concentration (Marchesiello et al., 2009).

The model bathymetry is derived from the Global Multi-Resolution Topography (GMRT) Synthesis (Ryan et al., 2009) smoothed with a maximum slope factor of 0.25 to reduce horizontal pressure gradient errors (Sikirić et al., 2009). The southern and eastern open boundaries are nudged to the six-hourly data from the Hybrid Coordinate Ocean Model - Navy Coupled Ocean Data Assimilation (HYCOM-NCODA) Analysis system. Six-hourly atmospheric forcing files (wind stresses, heat fluxes, and daily precipitation) are from the European Centre for Medium-Range Weather Forecast ERA-Interim reanalysis (Poli et al., 2010). Daily freshwater discharges for the five main rivers in the GoM (Mississippi, Atchafalaya, Colorado, Brazos, and Apalachicola) from the United State Geological Survey (USGS) are converted to an equivalent surface freshwater flux that decays away from the river mouths at a constant rate as in (Barkan et al., 2017). River momentum flux and tidal forcing are neglected in this work because of their weak influences on the deep-sea area which is the focus of this study (Bracco et al., 2019; Gouillon et al., 2010). Initial conditions are created by interpolating the field of HYCOM on September 31^th^ 2014 to the CROCO grid; the first 4 months of the simulation are discarded as spin-up. CROCO fields are saved every hour for offline particle tracking. At 1 km horizontal resolution, the use of hourly averaged velocity fields introduces only a small error in the tracer advection (Choi et al., 2017; Smith et al., 2011).

### Larvae tracking

Three release experiments with different tracking periods and different types of tracers are conducted in this work to investigate the connectivity pattern of *P. biscaya* in the northern GoM. Specifically, 4489 neutrally buoyant Lagrangian tracers (hereafter referred to as larval particles or particles) are deployed uniformly in 0.05 × 0.05º boxes centered at the location of known populations (Table S1, Figure 1a, red box with a colored text box next to it) and at 11 intermediate sites (green boxes) that could host *P. biscaya* populations according to habitat suitability modeling predictions (Georgian et al., 2020). These particles are tracked off-line (*release type 1 and 2*) using a Lagrangian tool developed to simulate ichthyoplankton dynamics (Ichthyop) (Lett et al., 2008) and are recorded hourly. Although the actual size of coral larvae is not infinitesimally small and could be slightly negatively buoyant (Brugler et al., 2013; Miller, 1998), the infinitesimally small approximation holds given the 1 km model resolution. A previous study has shown that in an environment with strong submesoscale features, slightly heavier/lighter (10%) buoyancy does not affect the transport significantly (Zhong et al., 2012). No other biological behaviors such as growth, mortality, settlement, and swimming are considered in this work, given that these are unconstrained for *P. biscaya*. The CROCO release depths are shown in Figure 1b. There is a 73 m difference on average between observations and model, and the largest discrepancy is found at GC852 (~ 200 m), where the observed bathymetry is very steep and varies greatly laterally on scales smaller than the model grid resolution.

A total of 76313 particles are released in the model layer above that at the seafloor on January 25^th^, April 25^th^, July 24^th^, and November 1^st^, 2015, and tracked for 56 days (release *type 1*). The pelagic larval duration (PLD) for *Paramuricea biscaya* is unknown, but Hilario et al. (2015) found that a PLD between 35 and 69 days may be representative of 50% to 75% of deep-sea species. In addition, the particles released at the 6 sampling locations on November 1^st^ are followed for another 92 days to evaluate connectivity over five months (~ 150 days in total, release *type 2*). Finally, the evolution of a dye released near the bottom (in the first s-layer) at the 6 sampling locations is simulated on-line (directly in CROCO) (release *type 3*) and followed for 120 days to explore the consistency between Lagrangian off-line and Eulerian on-line experiments.

### HYCOM hindcast

The inter-annual variability of the near-bottom currents is evaluated using the HYCOM hourly analysis at 1/25º horizontal resolution from 2010 to 2018 (Exp1, HYCOM/GOMl0.04) and the reanalysis data from 2010 to 2012 available at the same horizontal resolution but only at a three-hourly frequency (Exp2, HYCOM/GOMu0.04). The local circulations of these two experiments differ considerably over the common period because of the different choices regarding model configuration, vertical discretization and data assimilation routines (see https://www.hycom.org/hycom/documentation).

The velocity field is analyzed over the whole 2010-2018 period, and the dispersion patterns are simulated off-line using the HYCOM data in 2011, when both experiments are available and the derived near-bottom currents differ significantly between Exp1 and Exp2 and differ the most from those in 2015. Practically, by considering 2015 in CROCO and 2011 in the two HYCOM experiments we are exploring conditions as different as possible within the 2010-2018 period. Particle trajectories in HYCOM are advected using only the 2-dimensional near-bottom velocity field. To make the comparison with our simulations most relevant, we interpolated the original HYCOM velocity field at the horizontal resolution of 5 km and with the same vertical discretization used in the CROCO runs.

### Genetic data

To evaluate the performance of our models in predicting connectivity, we compared our results with the genetic connectivity estimates (migration rates *m*) by Galaska et al. (submitted). Briefly, Galaska et al. (submitted) produced single nucleotide polymorphisms (SNPs) data from individuals collected at the six main sites (DC673, MC344, MC297, MC294, GC852, and KC405) using the reduced representation DNA sequencing (RAD-seq) method (Baird et al., 2008; Reitzel et al., 2013). Migration rates (*m*), defined as the proportion of immigrant individuals in the last two generations, were estimated using BAYESASS v3.0.4.2 (Wilson and Rannala, 2003).

## Results and Discussion

### Circulation features in the GoM and its annual and inter-annual variability

Figure 2 shows the time-averaged near-bottom lateral velocities over the continental slope between 1000 and 3000 m, and their time-series where the coral sites are located, for CROCO in 2015 and for the HYCOM-NCODA analysis from 2010 to 2019. The spatial resolution difference among the two models implies that CROCO partially resolves submesoscale dynamics, while HYCOM-NCODA does not. The horizontal patterns of the averaged zonal current (west-east) of HYCOM outputs and our model result in 2015 (Figure 2a and 2c) illustrate the differences in current speed and, especially, variability (standard deviation) owing to CROCO’s higher resolution (Bracco et al., 2016). At the same time, some similarities are evident. For example, the prevalence of positive/negative velocities around (93.5°W, 27°N)/(93.5°W, 26.5°N), intermittent positive (eastward) and negative (westward) values between 94°W to 90°W, the presence of a westward velocity ‘belt’ following the 3000 m isobaths. The variability patterns between HYCOM and model outputs are also similar, just of stronger amplitude in CROCO, with the largest variability found between 94.5°W-93°W, near the Sigsbee Escarpment (89°W-91°W) and the Mississippi Fan (east of 90°W and south of 27°N). Energetic currents and large variability in the vicinity of the Sigsbee Escarpment are supported by field observations (Hamilton and Lugo-Fernandez, 2001). In the De Soto Canyon region (89°W-87°W, 27°N north), on the other hand, currents are weaker and less variable, as documented in previous studies (Bracco et al., 2016; Cardona et al., 2016).

**Figure 2.**
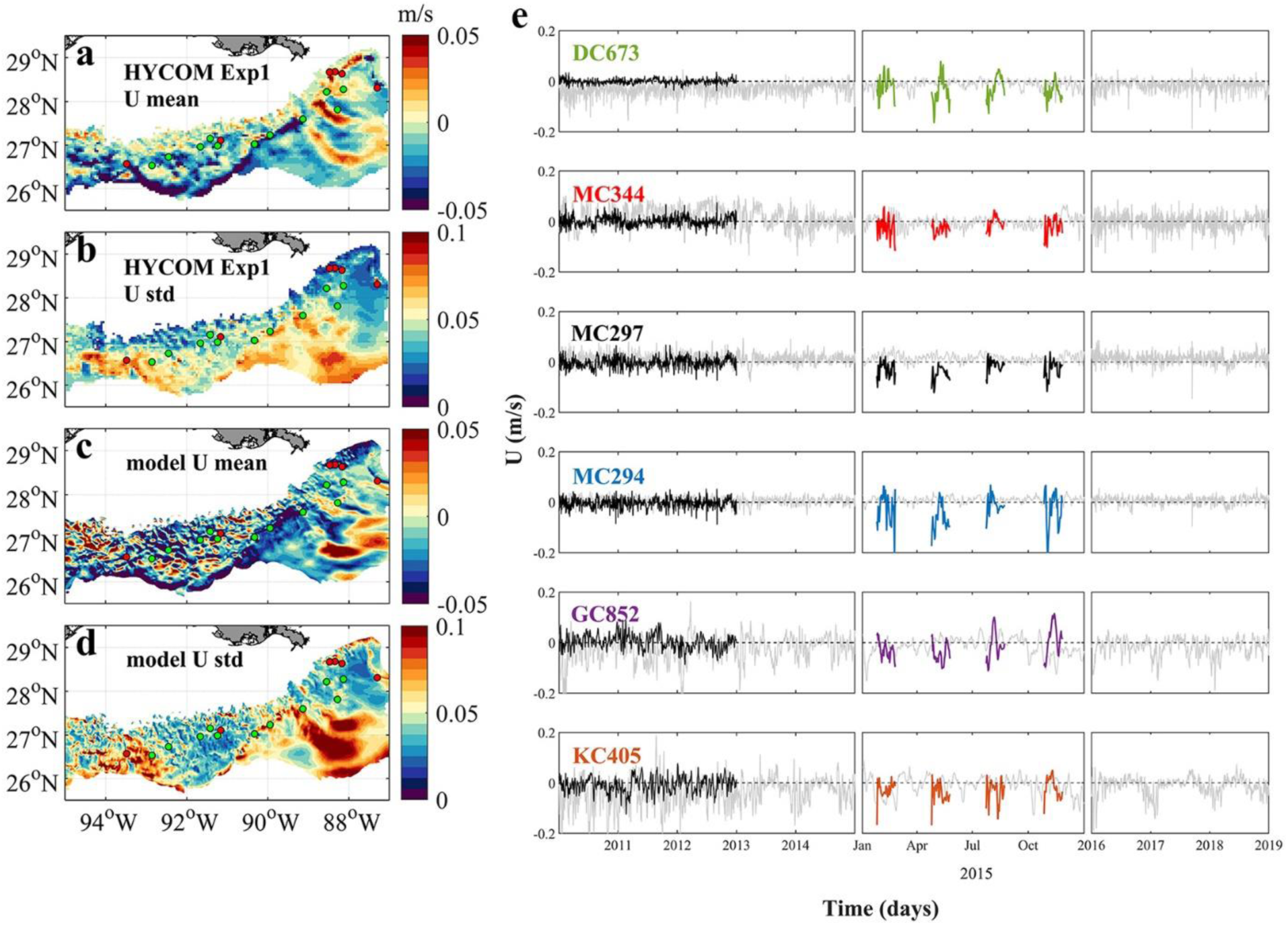
Near-bottom circulation in the northern GoM between 1000 m-3000 m. (a) averaged hourly HYCOM Exp1 zonal current (west–east) from 2010 – 2018, (b) the corresponding standard deviation (std), (c) daily averaged model zonal current, dataset is collected from February, May, August and November 2015 with a time period of 30 days for each simulation, and (d) the std of the modeled zonal current. The six sampling sites are colored by red dots, while the intermediate sites are shown in green. (e) shows the near-bottom zonal current at each sampling location during the period of 2010-2014 (left), 2015 (middle) and 2016-2018 (right). Grey, black and color lines indicate result of HYCOM Exp1, HYCOM Exp2 and CROCO, respectively. Positive values indicate an eastward current, and negative values westward.

Figure 2e depicts the time series of zonal near-bottom velocity at the six coral sites. In the period considered, there is some interannual variability but it is not much greater than across different seasons. Currents differ more for amplitude and direction between Exp1 and Exp2, than between Exp1 and CROCO. In 2015, the flow was westward (negative) in the monthly averaged HYCOM output (black thin line) and in the same direction, but generally stronger in CROCO. MC294 is the exception, and the directionality is reversed in HYCOM, but with a small amplitude. Overall in CROCO and Exp1 westward currents prevail around the coral sites.

Mesoscale and submesoscale circulations that may influence the transport of deep-water coral larvae can be seen in the normalized relative vorticity ζ */f* plots in Figure 3. ζ = *∂v/∂x - ∂u/∂y*, where *f* is the Coriolis parameter. *u* and *v* denote the zonal (west-east) and meridional (south-north) velocity components, respectively. *x* and *y* are the corresponding distances. The near-bottom vorticity indicates more intense submesoscale circulations over the continental slope in the western side of the domain compared to the eastern one, in agreement with previous work (Cardona et al., 2016). In 2015 slightly stronger submesoscale eddies are found in August compared to the other months, but this is likely the result of interactions between local currents and topography at that time, and is not indicative of a robust seasonal signal. Zoom-in fields of August and November results (Figure 3e and 3f) provide more details of the interactions between the near-bottom currents and topographic features. Overall, strong cyclonic submesoscale vortices (i.e., positive relative vorticity) are found in “valley” regions outlined by the close seafloor depth contours (see Figure 1a for better visualization) in both seasons. The formation of these small cyclones involves the shear layers and centrifugal instabilities associated with the mean flows and sloping boundary. The generation mechanism is beyond the scope of the present study but is discussed in previous works (Bracco et al., 2016; Gula et al., 2015; Molemaker et al., 2015). Away from these intense cyclonic vortices, GC852 locates in a less stable region with numerous weak, intermittent submesoscale structures.

**Figure 3.**
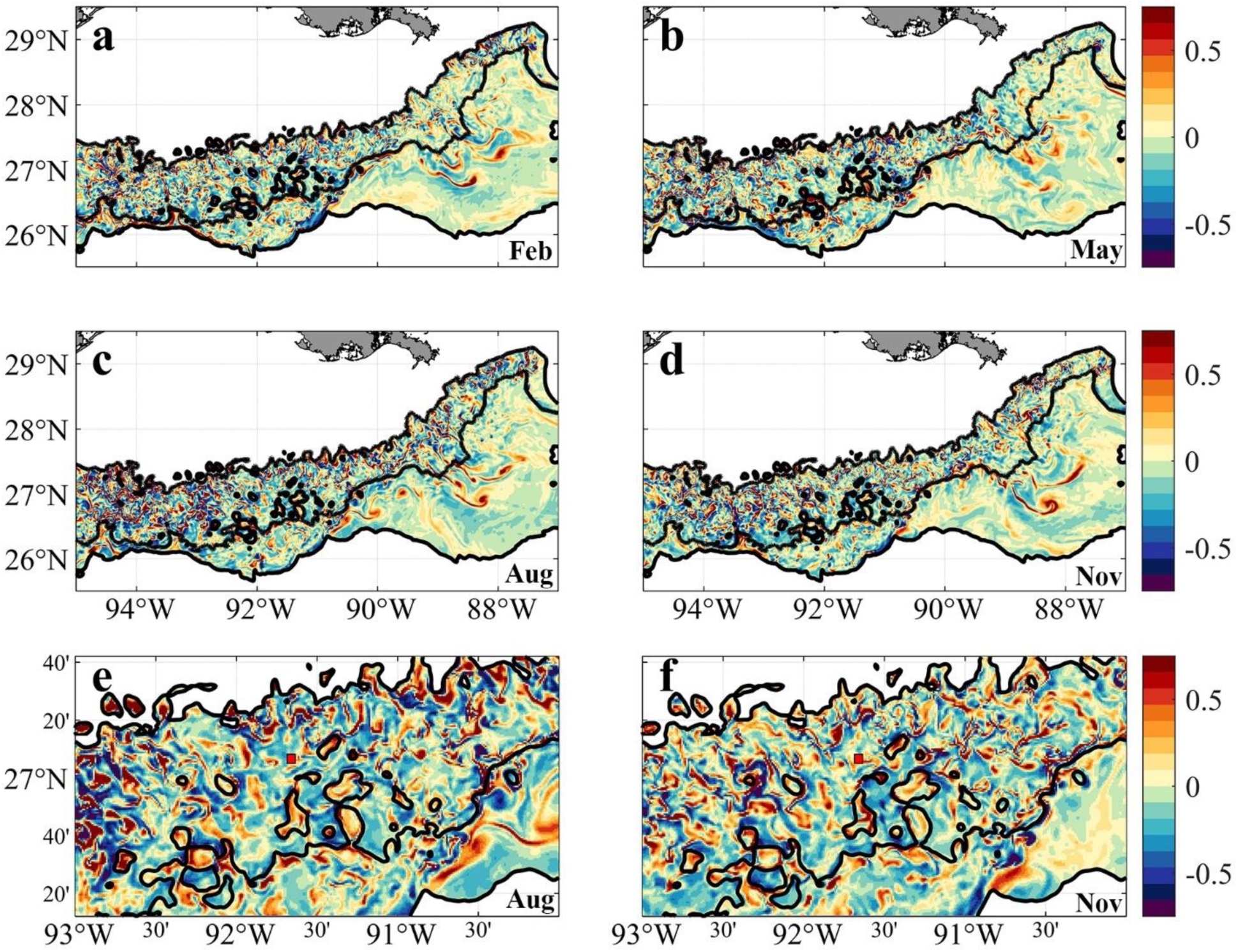
Relative vorticity near bottom in February (a), May (b), August (c) and November (d) calculated from model simulation. Negative values are anticyclonic or clockwise rotation. (e) and (f) are the zoom-in regions between 93-90° W, 26.2-27.7° N in August (c) and November (d). Red box with black edge shows the location of GC852.

### Genetic connectivity

The genetic analysis of the *Paramuricea* samples by Galaska et al (submitted) shows that migration rates are generally low (average *m* = 0.011), with a few exceptions (Figure 4). Considering the relative locations of sampled sites and the magnitude of the connectivity network, we can infer that the three sites in the Mississippi Canyon (i.e., MC344, 1853m; MC297, 1571m; and MC294, 1371m) are not well connected to each other (*m* < 1.5%). This is consistent with the depth-differentiation hypothesis that posits that genetic differentiation is greater across depth than geographic distance, even between sites relatively close by (Bracco et al., 2019; Quattrini et al., 2015; Galaska et al., submitted). Secondly, gene flow is predominantly westward. The DeSoto Canyon site DC673 is a source of genetic material to the Mississippi Canyon sites, particularly the deepest one (MC344). DC673 and KC405 (Keathley Canyon) appear well connected, despite been separated by 635 km of distance and 522 m of depth. These analyses also indicate that the Green Canyon site GC852 may also be an important source of immigrants to the Mississippi Canyon area because of eastward migration rates.

**Figure 4.**
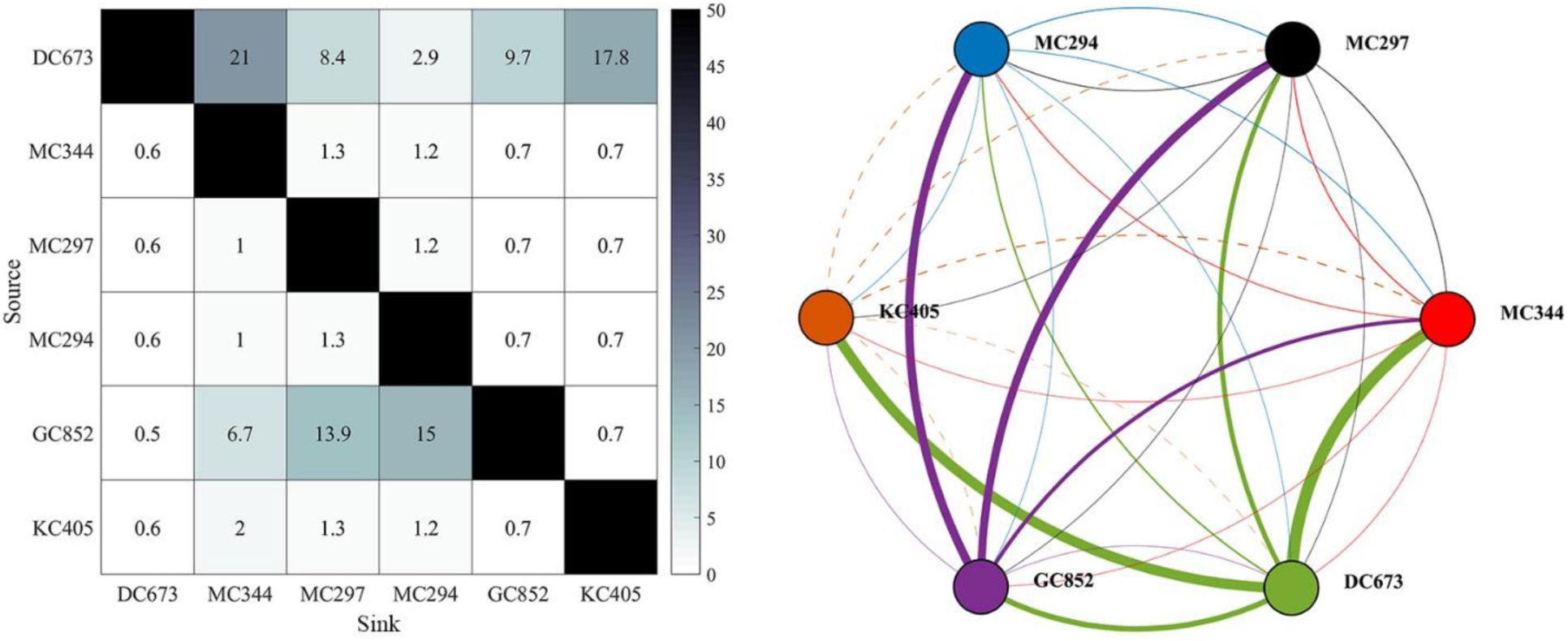
Connectivity matrix (left) and network (right) calculated from genetic data sampled at the six locations shown in Figure 1a. Figures are modified from Galaska et al (submitted). a) Matrix values correspond to point estimates of migration rates (*m*) as percentages (%). b) Network lines represent connections (dash line for KC405) and dots sites. Dots are color-coded by site. The color of the lines indicates the source site for the connections. Linewidths are proportional to *m* values. Sites are arranged from east to west (from DC673 to KC405).

### Connectivity pattern in the physical circulation model

The 2015 modeled connectivity patterns among the six sites are illustrated in Figure 5 where the horizontal and vertical distribution of Lagrangian particles is shown after 56 days in each season (Figure 5, *release type 1*). The horizontal dispersion of particles, even though characterized by detectable differences in the direction of motion and spreading area among sites, is mostly confined between the 1000 m – 2000 m isobaths in all seasons (Figure 5a-d). No obvious seasonal dependency is detected except for the relatively wider distribution of particles released from GC852 and KC405 in August, in response to the stronger submesoscale near-bottom flows along the western continental slope compared to the other months (see Figure 3e). In February, May, and November, most particles travel as far as 100 km to 300 km from their release locations, while in August particles released at GC852 and KC405 can be found as far as 600 km.

**Figure 5.**
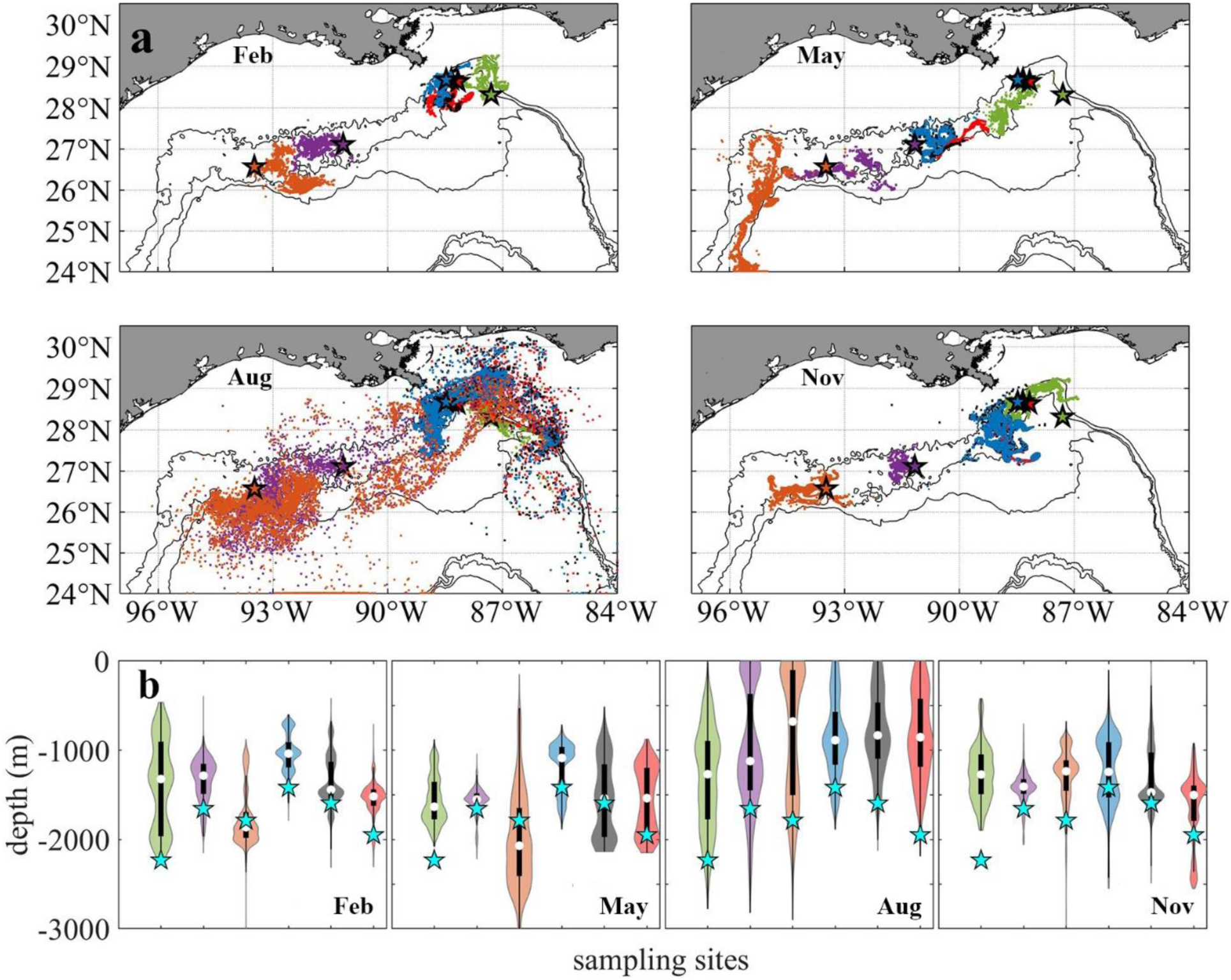
Distribution of Lagrangian larval particles released at the sampling sites after 56 days. Horizontal (a). Vertical (b). Particles from different sites in (a) are colored consistently with previous figures. For each violin in (b) the central white square indicates the median, and the bottom and top edges of the black box indicate the interquartile range between 25^th^ and 75^th^ percentiles, respectively. Thin black line shows 95% confidence level, and cyan stars indicate the initial release depth for each site. The width of each violin represents frequency, i.e., density plot.

As mentioned earlier, the near-bottom circulation at the GoM continental slope is predominately along depth contours, therefore larvae migration and connectivity are closely linked to the alongshore (lateral) direction of motion. Virtual larvae released at DC673, MC344, MC297, and MC294 move principally westward in all seasons, as also reported in Cardona et al. (2016). For KC405 and especially GC852 particles, on the other hand, eastward movement can be observed as their lateral velocity is more variable (Figure 2), especially for the February and August releases in the CROCO run. The greater variability of the circulation in the central portion of the GoM continental slope results from the many recirculation zones that occupy this area (see Bracco et al., 2016, their Figures 3 and 10).

In the vertical direction (Figure 5e), the particle spreading is highly variable but without a seasonal trend. For the August release, a large portion of particles that originated at GC852 and KC405 are displaced by more than 500 m in two months. DC673 is a site characterized by strong eastward and westward currents and high variance in particle displacement (Figure 2a-d) likely caused by the steep slopes of the surrounding topography (Figure 1a). On the contrary, particles released at the Mississippi Canyon sites in all seasons, and GC852 in February, May, and November show smaller variances in their trajectories.

The horizontal dispersal patterns for 2015 are quantified by the 2-dimensional kernel density estimation (KDE) using all four releases and the November extended one as well (Figure 6). The KDE is a non-parametric technique to produce a smooth probability density function given a random variable. The figure shows the heat map of particles’ trajectories in the first 10, 30, 56 days and all available releases, i.e., 148 days in November and 56 days in all the other seasons. Particles displaced in the vertical direction by more than 800 m depth are discarded (~ 20% of all particles and mostly ending in very deep water). For all cases, high KDE values are found in regions within ~ 100 km from the release points. A clear pathway from the northeast region near the De Soto Canyon to the southwest area between KC405 and GC852 is outlined following the 2000 m isobath. In addition to the prevailing westward transport, an eastward branch stemming from KC405 could be responsible for completing the east-to-west connectivity.

**Figure 6.**
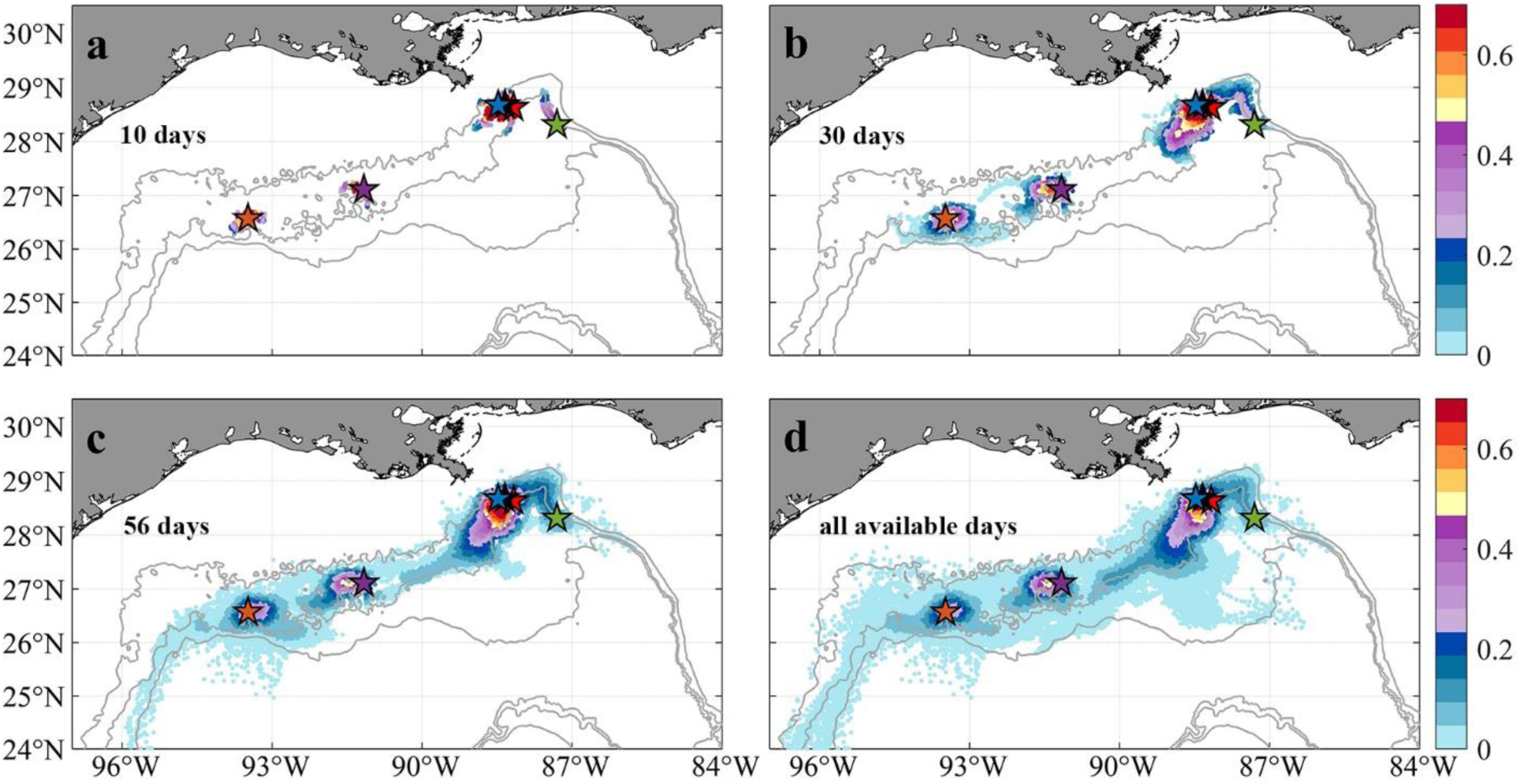
Horizontal probability distribution of Lagrangian larval particles after their release at the six sampling sites (colors indicate probability values). Maps of the kernel density estimation (KDE) of particles passages during periods of 10 days (a), 30 days (b), 56 days (c) and all available days (d, 56 days in February, May and August, and 148 days in November) since release. Subsampling (sample every 20 particles spatially and 2 days temporally) has been applied for plotting. Few larval particles that are displaced more than 800 m in the vertical direction away from the bottom are removed to focus on near-bottom processes.

The potential connectivity network from the model integrations is compared to the observed one from the genetic data in Figure 7. Even though the extended tracking period in November leads to more horizontal (increased from 5 to 6 connections) and vertical (from 2 to 4 connections) connections among the six sampling sites, the modeled network is still less dense than the measured one (genetic), possibly indicating an overall underestimation of coral connectivity. When the vertical dimension is considered, the connectivity from KC405 to GC852 is lost (dash line) and the probability values of several other connections decrease significantly. In line with the genetic evidence, the three MC sites are all connected horizontally, but their connectivity decreases by 64.6% when depth is considered. Connections between the easternmost DC673 and the three MC sites are also observed in the modeled network and are most evident for the DC673-MC344 pair. The model, however, fails to simulate both the long-distance bi-directional communication between DC673 and KC405 and the connectivity out of GC852. In other words, KC405 and GC852 are statistically isolated from the eastern sites in the 2015 CROCO simulation, contrary to the outcome from the genetic inferences.

**Figure 7.**
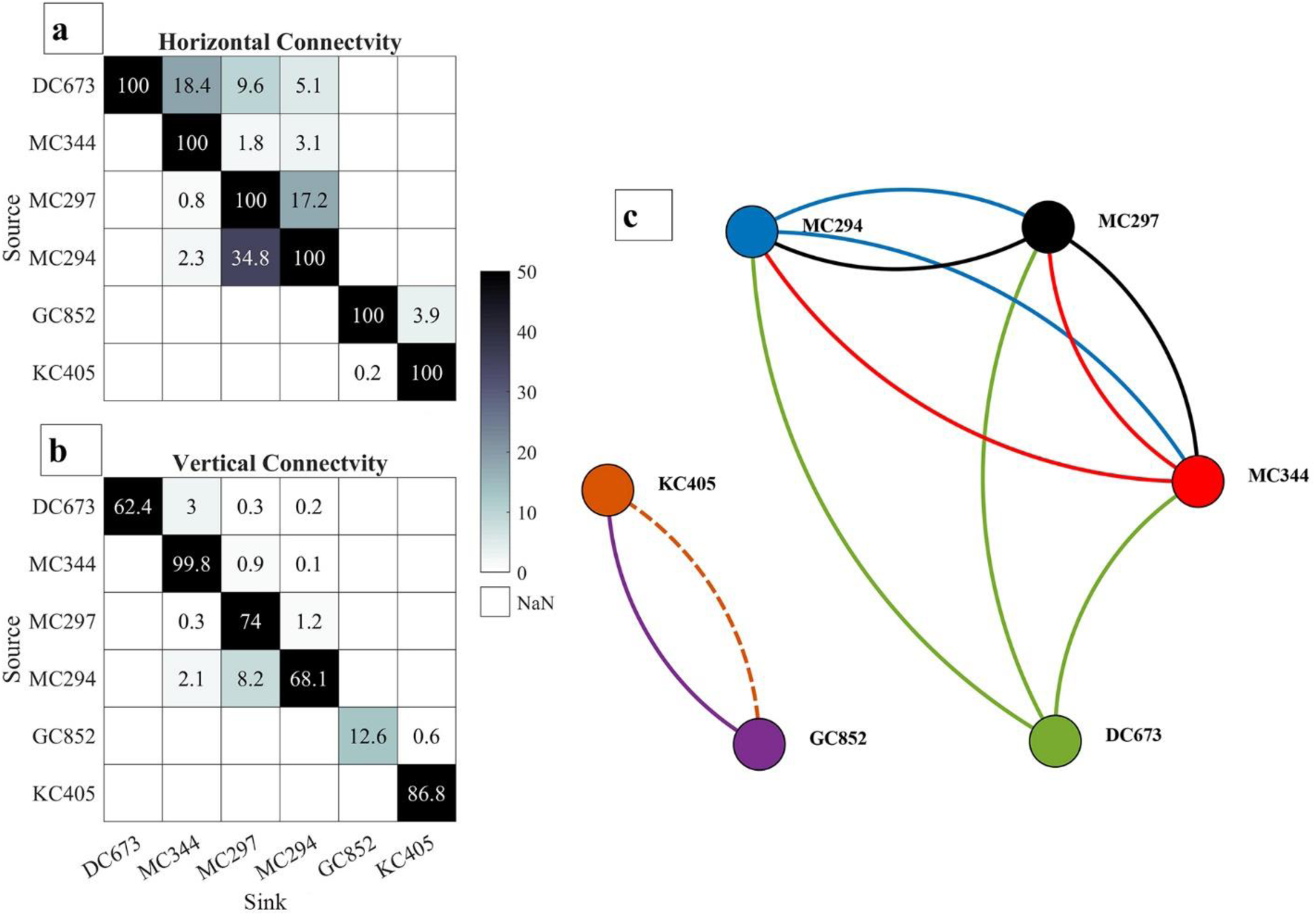
Horizontal *l*_*h*_ (a) and 3D *l*_*v*_ (b) connectivity probability (%) matrices and connectivity network (c) calculated from the Lagrangian simulations performed over 56 days in February, May and August, and 148 days in November. Line colors indicates the source site for each connection. Solid lines indicate three-dimensional connectivity and the dash line represents horizontal connection only. Line width does not correspond to the magnitude of connectivity. Sites are arranged from east to west (from DC673 to KC405).

### The potential role of intermediate populations

The six sampled sites do not represent the only sites hosting *P. biscaya* populations in the northern GoM. A habitat suitability model recently published by Georgian et al. (2020) predicted a broad distribution of habitat areas where *P. biscaya* is likely present. Using this model, we selected eleven other potential sites distributed between 1,600 and 2,300 meters deep, and roughly equidistantly between the six sampled sites (distances among all sites ranging between approximately 50 and 100 km, Figure 1a). We integrated larvae trajectories from these sites and investigated potential connectivity, under the assumption that these locations are indeed populated by *P. biscaya* and can thus participate in larval exchange. The connectivity matrices in Figure 8 show the probabilities of the larval exchange among all sites (sampled and predicted). These results clearly show that predicted connectivity is predominantly westward across the region. The probability of connectivity decreases as a function of distance, but depth ultimately dictates whether or not neighboring sites are connected. This is because the connectivity among sites that are relatively close, geographically, is limited by diapycnal mixing across depth.

**Figure 8.**
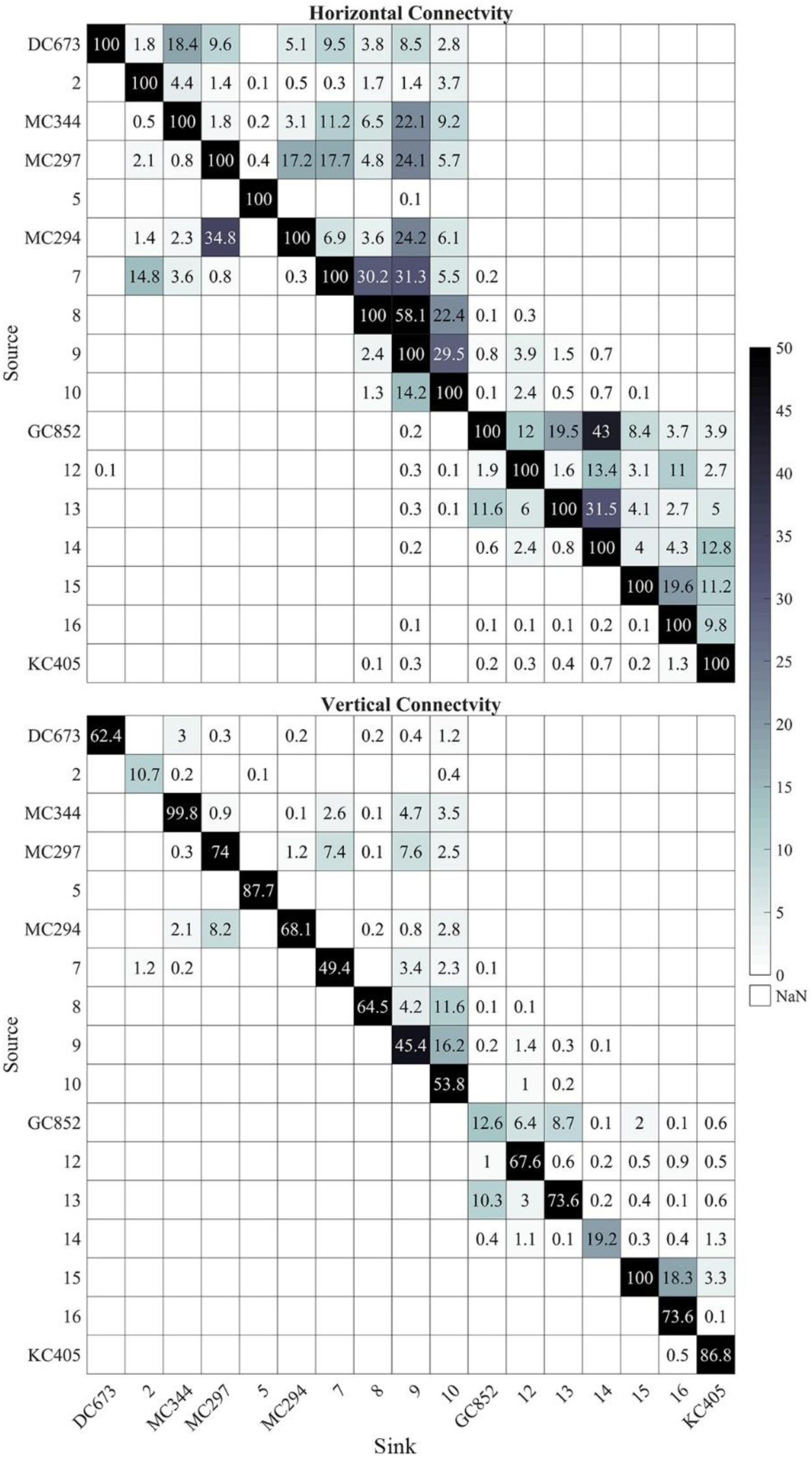
Heat maps of horizontal *l*_*h*_ (a) and 3D *l*_*v*_ (b) connectivity probability (%) calculated from all 17 sites (six sampled and eleven predicted). The connectivity is calculated from 56 days output in February, May and August and 148 days output in November, 2015. Connectivity values below 0.1% are not shown. Sites are arranged from east to west (from DC673 to KC405).

We built a probabilistic graphic model to quantify the role of intermediate sites in metapopulation connectivity and visualize the modified connectivity matrices. We adopted a directed cyclic graph, instead of the most commonly used Bayesian network, which is a directed acyclic graph (Ben-Gal, 2007). Based on the concepts of conditional probability and chain rule, the joint probability of three events from A (source) to B (intermediate site) to C (sink) can be represented as P(A, B, C) = P(A)×P(B|A)×P(C|A, B). The cyclic network built here allows us to understand the dependency among events (nodes) and assigns probabilities (edges) to them. Our goal was to identify as many connections as possible using the current matrix rather than quantifying the probability/strength of each connection after certain iterations, therefore connections (graph edges) that occur more than once are excluded for the following iterations.

Figure 9 presents the 3-dimensional connectivity pattern and network (based on the vertical matrix in Figure 8) after three iterations/steps (no new connections are found after four or more iterations). Long-distance east to west connections are resolved after three steps (two intermediate sites acting as stepping stones) for particles released from MC344 (Figure 9a). Meanwhile, GC852 particles show both eastward and westward predicted connections despite the direct exchange terminates at an intermediate site around 89°W (Figure 9b). The complete connectivity network (Figure 9c) resolves 19 out of the total 30 possible connections and provides a mechanism for long-distance connectivity between MC344 and DC673 through stepping-stone dispersal mediated by intermediate sites. However, the inferred eastward gene flow from GC852 to the Mississippi Canyon sites remains unexplained by the model.

**Figure 9.**
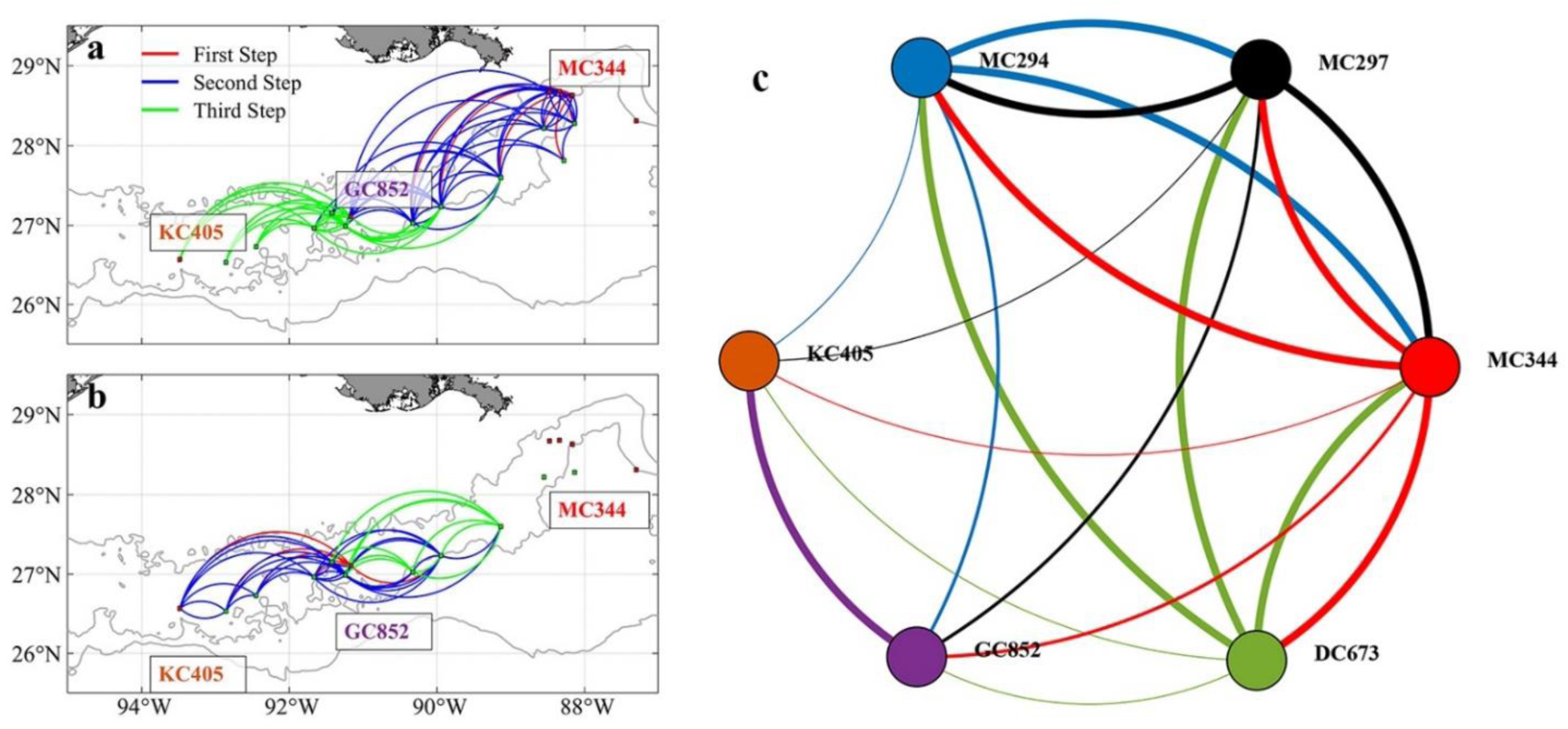
Predicted 3D patterns of metapopulation connectivity considering the suitable intermediate sites. Connections found after three steps for larval particles released from sites MC344 (a) and GC852 (b). Line colors in (a) and (b) indicate the step at which a connection between two sites was found. Connectivity network among the six sampled sites calculated after including intermediate sites (c). Line colors in (c) indicate the source site for each connection. Linewidth in (c) indicates direct (thick lines, first step) or mediated connections (regular and thin lines, second and third steps, respectively).

### Connectivity with extra vertical diffusion

In the work presented so far, vertical and horizontal diffusivities are parameterized at the CROCO grid size. In CROCO, the modeled *k*_*z*_ is comparable to observed values (1.3×10^−4^ - 4×10^−4^ m^2^s^-1^) calculated from a dye injection experiment conducted in 2012 at the Deep-water Horizon site (Ledwell et al., 2016) (see e.g. Figure A1 in Bracco et al., (2019)). Following previous studies, an additional vertical diffusion coefficient *k*_*z*_ of 10^−4^ m^2^s^-1^ is introduced to the tracking model to recognize the uncertainty associated with larval buoyancy and a likely underestimation of diapycnal mixing very close to the bottom at the continental slope. Since we are exploring the potential impacts of extra perturbation other than simulating the actual marine environment where *k*_*z*_ varies spatially (Visser, 1997), a naïve random walk model with constant diffusivity (Hunter et al., 1993) is adopted to track larval particles released in November 2015 for 148 days (Figure 10). Both horizontal KDE and vertical probability density function (PDF) are nearly indistinguishable from the ones obtained without added vertical diffusivity after five months. The additional vertical diffusion increases only slightly (by 0.1%) the chances of horizontal connectivity from MC297 to MC344 (not shown).

**Figure 10.**
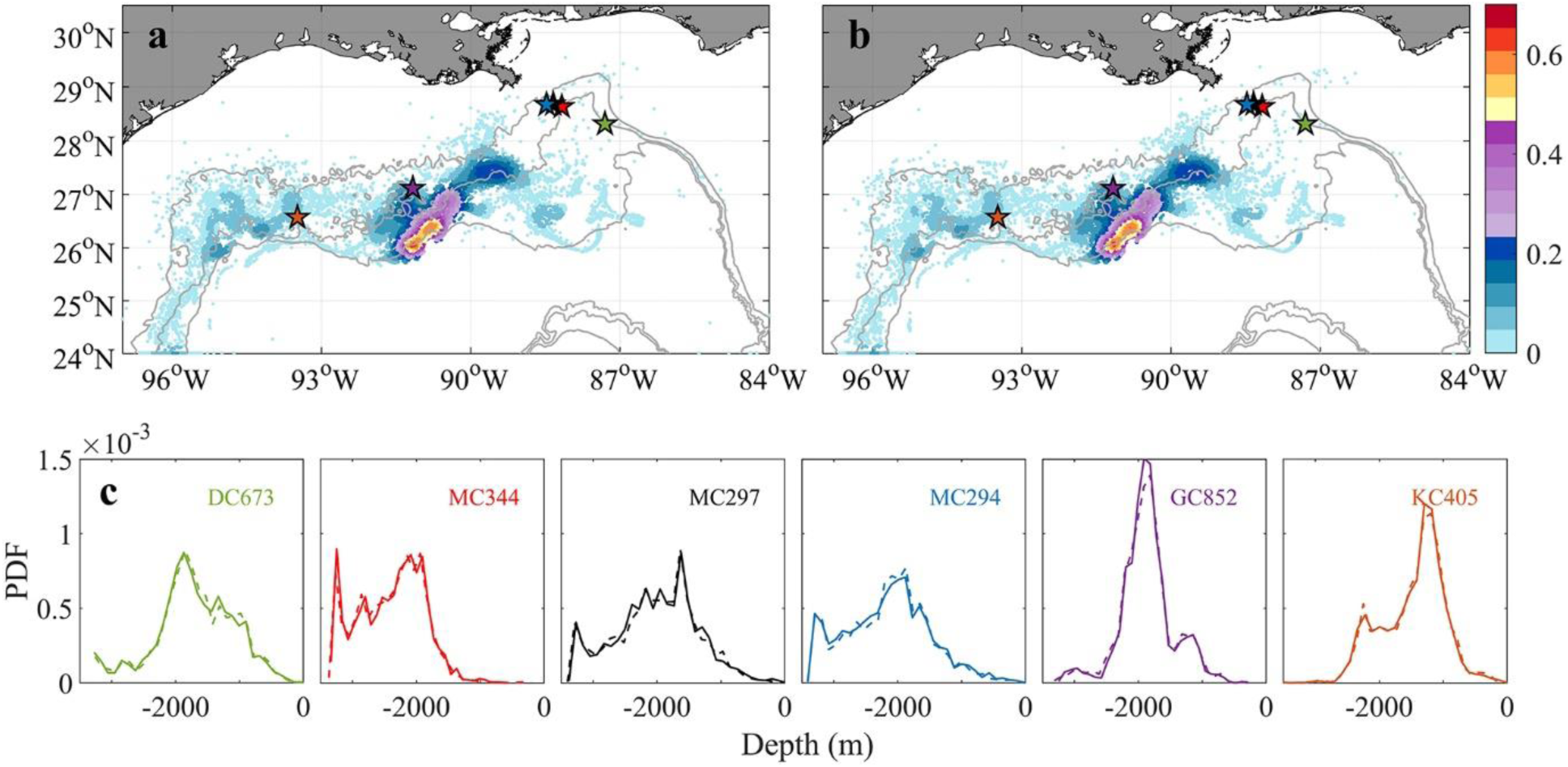
Horizontal kernel density estimation (KDE) pattern of particle locations after 148 days tracking in November without (a) and with (b) extra vertical diffusion (colors indicate probability values). A comparison of the probability density function (PDF) of the depth of particles released at each location is shown in (c). Solid/dash line indicates without/with vertical diffusion. All available particles are retained to show a thorough comparison.

### Lagrangian and Eulerian tracking comparison

To further validate the off-line Lagrangian larval particle integrations, we released a dye on-line in the bottom model layer at the six sites in a 0.05 × 0.05º box in November and tracked it for 120 days (*release type 3*). Figure 11 compares the Lagrangian and Eulerian dispersion based on the distribution of larval particles and the absence/presence of connectivity among sites (Table 1). We stress that the release depths for the Lagrangian larval particles and the dye differ, and that diffusion is included in the dye momentum equations, while this is not the case for the Lagrangian particles (see e.g., Bracco et al., 2009 and more recently Paparella and Vichi, 2020 for pros and cons of using Eulerian versus Lagrangian approaches). Figure 11 shows the dye concentration field with superimposed particle positions from *release type 2* after 120 days of simulation. The results demonstrate that although Lagrangian particles cover a smaller area, they capture the main dispersal features successfully in most cases, e.g., MC344, MC297, GC852, and KC405. For DC673 and MC294, the notable differences are due to the fact that particle positions are plotted at a specific time step (after 120 days), rather than displaying the overall trajectories that would better match the dye pattern. In the end, when comparing connectivity using larval particle trajectories and the 3D dye distribution, it is apparent that the first provides a lower bound to both the horizontal and vertical dispersal due to the lack of subgrid diffusion and not perfectly resolved vertical velocities when using hourly averaged outputs (Wagner et al., 2019).

**Table 1.** Difference in Eulerian and Lagrangian connectivity results, H or V indicates that horizontal or vertical connectivity exists in Eulerian but not in Lagrangian result, while HV indicates both horizontal and vertical connectivity are observed in Eulerian but not in Lagrangian result. Note blank cell means Eulerian and Lagrangian return same predictions (either connections exist or not exist).

| Sink | DC673 | MC344 | MC297 | MC294 | GC852 | KC405 |
| --- | --- | --- | --- | --- | --- | --- |
| Source |  |  |  |  |  |  |
| DC673 | - |  |  |  |  |  |
| MC344 | H | - | H | H | HV |  |
| MC297 | H | H | - |  | HV |  |
| MC294 | HV | HV |  | - | HV |  |
| GC852 |  | HV | H | H | - | V |
| KC405 |  |  |  |  | HV | - |

**Figure 11.**
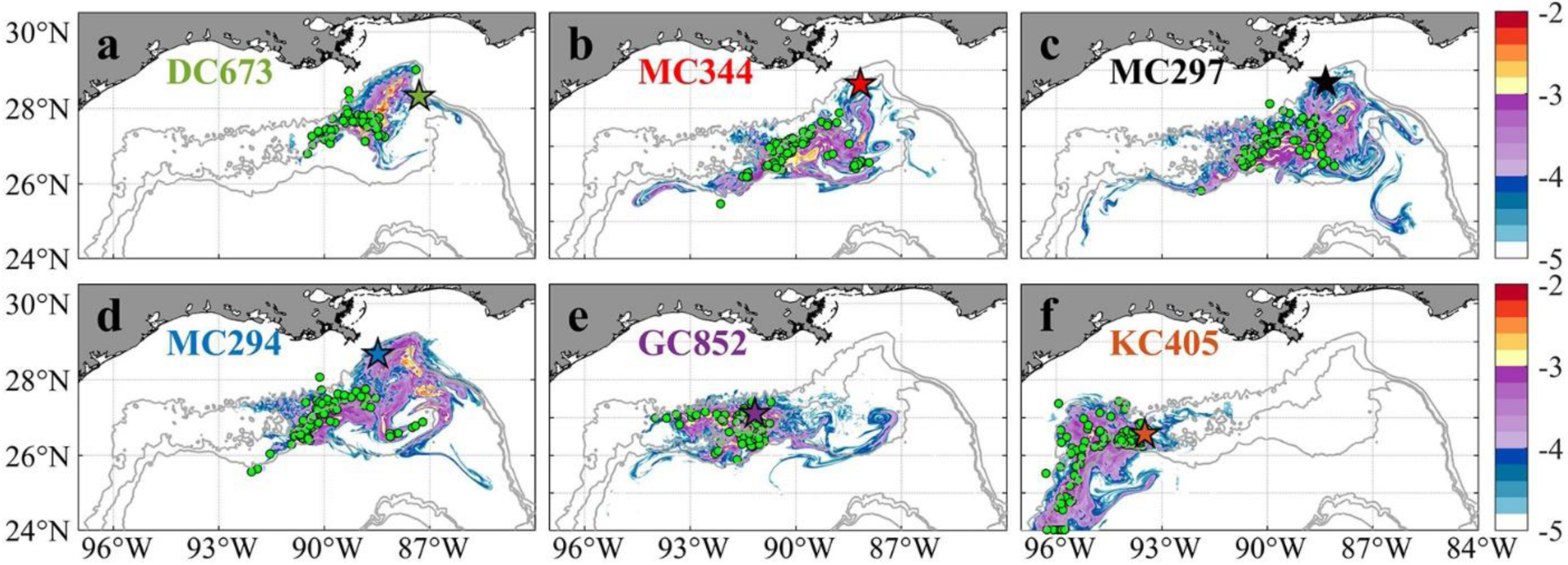
Horizontal distribution of integrated Eulerian dye concentration normalized by initial tracer concentration (log_10_ scale, same as Bracco et al., 2018) and Lagrangian larval particles after 120 days from their release on November 1^st^, 2015 for sampling site at DC673 (a), MC344 (b), MC297 (c), MC294 (d), GC852 (e), and KC405 (f). Colored pentagram with black edge shows release location. Green circles indicate the positions of Lagrangian particles on the same day. Again, particles displaced vertically more than 800 m from the bottom are not included in the calculation to focus on near bottom processes. The number of particles is randomly subsampled for visualization purposes.

### Inter-annual variability investigation using ocean hindcast data

Even in the Eulerian dye approach, the direct connections from GC852 to the Mississippi Canyon site inferred from the genetic data are missing because of the prevailing westward near-bottom current in 2015. While the submesoscale permitting resolution allows for a better representation of the bathymetry and circulation, it increases the computational time and limits the time period of the exploration. To partially address this issue, we examine the *Paramuricea* connectivity in 2011 when a more persistent eastward circulation was found in the HYCOM hindcast data, especially in Exp2. Using only the near bottom horizontal velocity components for the advection, we released the same number of larval particles as done for CROCO on April 1^st^ 2011, when the mean currents are eastward along the 1000 – 2000 m slopes, and follow them for 148 days. Given the lower frequency at which velocities are saved in Exp2 (3 hourly only), and the underestimation of mean velocities and their standard deviations compared to CROCO results, we also attempted to track the particles by using twice the velocities values reported in the hindcast. Figure 12 provides the horizontal distribution of larval particles after 56-day and 148-day tracking for Exp1 and Exp2 horizontal velocities. The prevalent spreading direction for the particles remained westward, as in CROCO. Greater spreading is found in Exp1 compared to Exp2, and patterns are significantly different. We speculated that the choice of vertical discretization, with better near-bottom vertical resolution in Exp1, explains the differences. Results from Exp1 compare well in terms of overall connectivity patterns with those of CROCO, despite the different times considered (2011 in HYCOM Exp1 and 2015 in CROCO).

**Figure 12.**
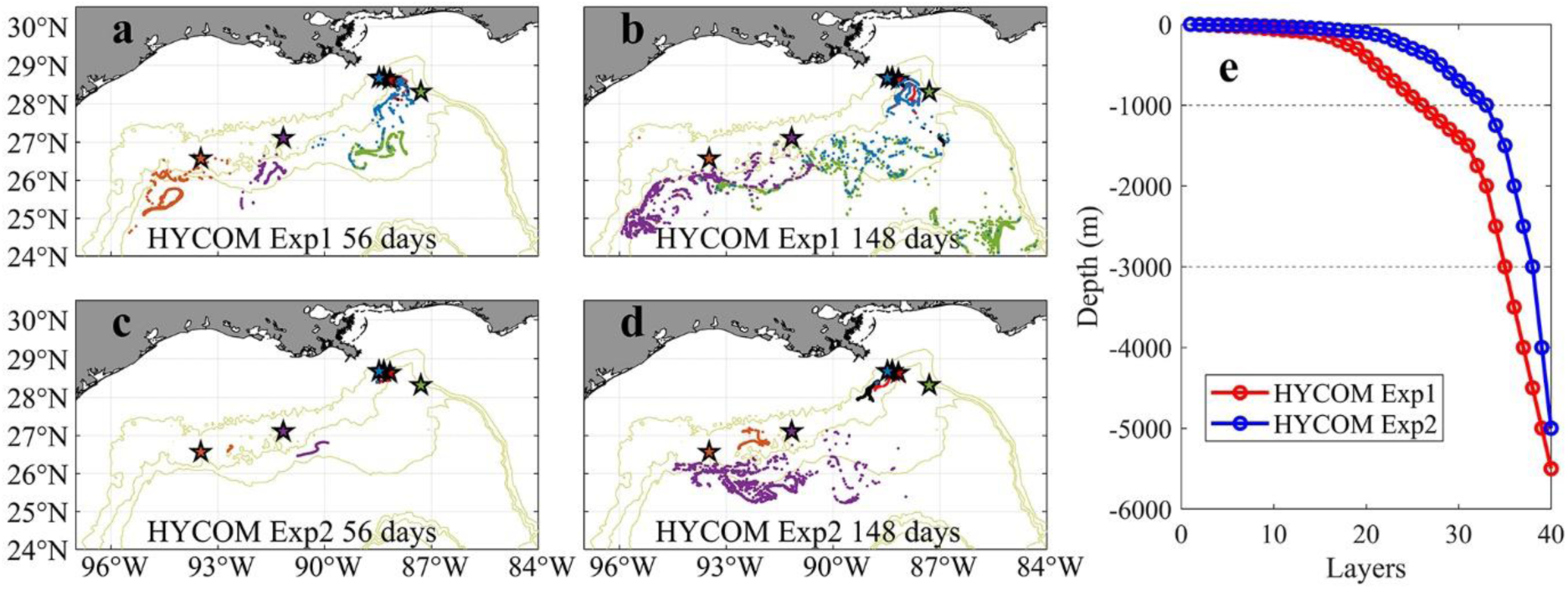
Near-bottom distribution of Lagrangian larval particles released in 2-dimensional HYCOM Exp1 (a, b) and HYCOM Exp2 (c, d) fields in April 2011 after 56 days (a, c) and 148 days (b, d) tracking, respectively. Each dot represents the position of a particle. Particles from different sites are colored consistently with previous figures. The setup of vertical layers in each experiment is shown in (e).

The traveled distance of the larval particles released at each site in 2015 (CROCO) and 2011 (HYCOM) is shown quantitatively in Figure 13, where the outcome for doubling the horizontal velocities in Exp2 is also displayed. As to be expected, given our selection of a year with conspicuously different currents, inter-annual variability is observed.

**Figure 13.**
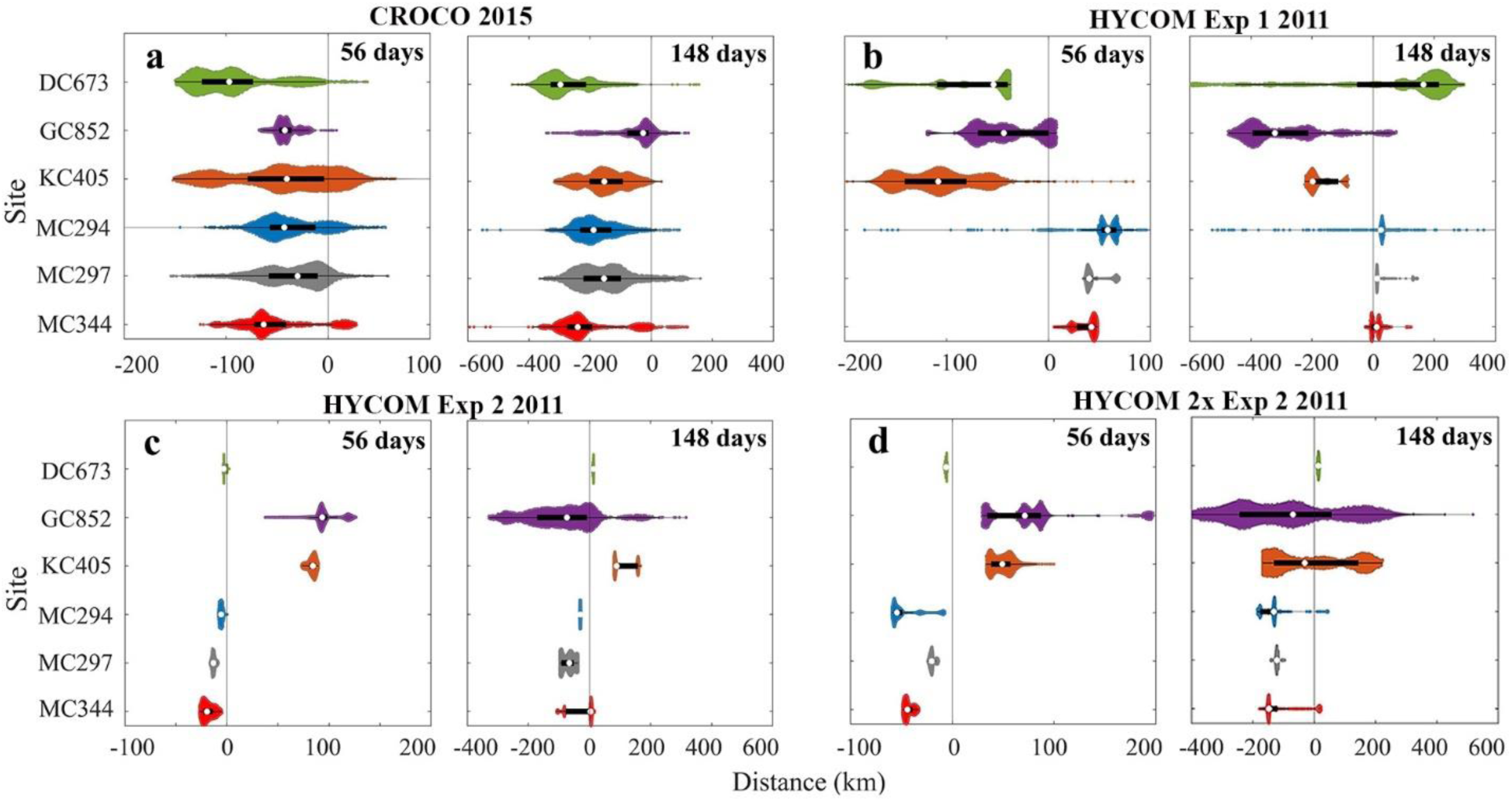
Distribution of horizontal dispersal distances for larval particles after 56 and 148 days. November 2015 release using CROCO (a), and April 2011 release using HYCOM for Exp1 (b), Exp2 (c) and Exp2 with doubled zonal velocity (d). Positive values represent eastward movement and negative westward. Particles with a vertical displacement larger than 800 m are removed in CROCO result. The definition of color and violin plot is consistent with Figure 5.

The 75^th^ percentiles of the distance traveled by KC405 particles in the 2011 HYCOM simulation and DC673 particles in the 2015 CROCO run are ~ 180 km and ~ 300 km (148 days), respectively. These numbers suggest that at least a 10-18 months pelagic larval duration (PLD) is needed to achieve long-distance direct connectivity between KC405 and DC673 if no intermediate sites are considered and without including the vertical aspect. Westward spreading of larval particles in both 2011 and 2015, the generally westward currents in the nine years considered for HYCOM Exp1, and the weak standard deviation detected in Figure 2 suggest the predominance of an east-to-west along-isobath pathway of dispersal for the three Mississippi Canyon sites. Finally, it is worth noting that by examining the difference between 56-day and 148-day results, a fraction of the released particles, e.g., at GC852 (~ 17%) in 2015, and at MC344 (~ 22-61%) and DC673 (~ 70-89%) in 2011, change their traveling directions with respect to their initial locations, indicating a role of seasonal variability.

### Summary and Conclusions

In this work, an integrated larval dispersal framework consisting of a high-resolution regional hydrodynamic model (ROMS-CROCO) and a Lagrangian larval particle tracking model (Ichthyop) were performed to predict the dispersal patterns and potential metapopulation connectivity of *Paramuricea biscaya* in the northern GoM. Lagrangian deployments with 76313 larval particles were conducted in different seasons for up to ~ 150 days and validated by a comparable Eulerian dye experiment in November. The potential contributions of vertical diffusion and intermediate sites to larval connectivity were also studied. The role of inter-annual variability of near-bottom circulations was investigated using HYCOM hindcast data.

The output of our biophysical model showed a mostly congruent agreement with the estimated genetic connectivity patterns (Galaska et al. submitted). In CROCO we found a prevailing westward pathway following the ~ 1000 – 2000 m isobath along the continental slope of the northern GoM regardless of seasons in 2015. In general, our estimations of dispersal distances (less than 100 km in 56 days to 300 km in 148 days) agreed well with previous deep-sea studies that considered pelagic larvae duration from 40 days to 1 year (Breusing et al., 2016; Cardona et al., 2016). Strong horizontal but significantly reduced vertical, and therefore 3-dimensional, connectivity among sites near the De Soto Canyon (i.e., DC673, MC344, MC297, and MC294) further confirms the depth differentiation hypothesis in agreement with previous studies (Bracco et al., 2019; Quattrini et al., 2015). The predominantly westward currents and weak variance near the 1000 m isobath in the De Soto Canyon region found both in HYCOM and in the CROCO model explain the westward confined pathway along the geographic feature shown in Figure 6. In contrast to the relatively stable hydrodynamic environment around the MC sites (also reported in Bracco et al. (2016)), highly variable currents over complex topography occupy the central and western portion of the continental slope, around GC852 and KC405 (Figure 1a and Figure 2). Such variability can result at times in eastward transport, as verified in CROCO in February and August 2015 (Figure 5 and Figure 6) and in HYCOM in April 2011, with high variance in the modeled displacement of larval particles in both models, and contributes to the diversity of connectivity patterns found in this region.

The inter-annual variability in the near-bottom circulation may be responsible for the few incongruences with the genetic connectivity estimates, for example, the fact that eastward gene flows from GC852 and KC405 is underestimated by the model. Inter-annual variability in the study region has been partially documented by Cardona et al. (2016) in their 3-year simulations from 2010 to 2012. In agreement with our 2011 HYCOM findings, they found several eastward transport events of larval particles originating at ~ 92Wº to the central Mississippi Canyon (~ 89Wº) and a predominant westward transport between 600 and 1000 meters depth near the De Soto Canyon.

The long-distance genetic connectivity between DC673 and KC405 may be explained by direct dispersal if we assume a pelagic larval duration of at least one year for *Paramuricea biscaya*. However, this possibility seems unlikely. Pelagic larval durations (PLD) of more than a year have been documented for a few deep-sea invertebrate species (Young et al. 2012), but not for deep-sea corals. Shorter PLDs, between 35 and 69 days, may be representative for most deep-sea species (Hilario et al. 2015). A probabilistic graphic model suggests that stepping-stone dispersal mediated by intermediate sites provides a more likely mechanism for long-distance connectivity between the populations in De Soto and Keathley canyons.

We briefly compared Lagrangian and Eulerian approaches in estimating larval dispersal patterns (Figure 11 and Table 1). Our results implied that the Lagrangian larval particle trajectories computed by interpolating the model’s hourly averaged velocity underestimate vertical velocity and sub-grid diffusion compared to the Eulerian approach (Ali Muttaqi Shah et al., 2017; Wagner et al., 2019). The Lagrangian-derived connectivity represents therefore a lower bound of the Eulerian one. The Lagrangian method presents, on the other hand, several advantages, providing the opportunity to perform multiple sensitivity integrations off-line, and add at low computation cost biological and behavioral constraints, such as mortality, swimming, growing, and settlement. The choice of approach should be done considering the application domain and question(s) in hand.

Our results emphasize the need for multi-year simulations, or at least multi-year analyses of the velocity field, to quantify dispersal patterns in the deep ocean, especially for bi-directional and long-distance connectivity. It is also known that depth and topographic slope are key factors determining the suitability of a habitat for many deep-water corals (Hu et al., 2020; Kinlan et al., 2020; Georgian et al. 2020). Thus, detailed topographic mapping, high horizontal resolution and the fine-scale vertical resolution near the ocean bottom should be adopted to reduce uncertainty in the model representation of bottom currents. Finally, in this work we focused on dispersal processes, but larval traits, e.g., swimming, settlement, and mortality, should be further investigated to improve the realism of modeling studies of coral connectivity.

## Acknowledgement

Funding support for this project was provided by the National Oceanic and Atmospheric Administration’s RESTORE Science Program award NA17NOS4510096 to Lehigh University (Herrera, Bracco and Quattrini co-PIs). Sampling was supplemented by previous sampling efforts under the Lophelia II Project funded by BOEM and NOAA-OER (BOEM contract #M08PC20038) led by TDI-Brooks International, the NSF RAPID Program (Award #1045079), the NOAA Damage Assessment, Response, and Restoration Program, and ECOGIG (Gulf of Mexico Research Initiative). We thank Chuck Fisher, Erik Cordes, Illiana Baums for leading those supplemental field efforts and providing access to samples and dive time. We thank Cathy McFadden, Cheryl Morrison, and Frank Parker for project support.

