## Supplemental Table 1 for "Kilometer-scale larval dispersal processes predict metapopulation connectivity pathways for *Paramuricea biscaya* in the northern Gulf of Mexico"

### Supplementary

Table S1. Sampling site location and depth information. - = intermediate sites as predicted in Georgian et al. 2020

| Site Id | Site Name | longitude | latitude | Depth (m) |
|---|---|---|---|---|
| 1 | DC673 | -87.3015 | 28.3124 | 2222 |
| 2 | - | -88.1365 | 28.2794 | 2143 |
| 3 | MC344 | -88.1695 | 28.6335 | 1853 |
| 4 | MC297 | -88.3423 | 28.6799 | 1571 |
| 5 | - | -88.2808 | 27.8141 | 2219 |
| 6 | MC294 | -88.4765 | 28.6722 | 1371 |
| 7 | - | -88.5503 | 28.2196 | 1708 |
| 8 | - | -89.1350 | 27.6003 | 1622 |
| 9 | - | -89.9449 | 27.2348 | 1936 |
| 10 | - | -90.3286 | 27.0269 | 1885 |
| 11 | GC852 | -91.1651 | 27.1106 | 1410 |
| 12 | - | -91.2405 | 26.9883 | 1967 |
| 13 | - | -91.4224 | 27.1544 | 1915 |
| 14 | - | -91.6654 | 26.9647 | 1918 |
| 15 | - | -92.4495 | 26.7302 | 1667 |
| 16 | - | -92.8585 | 26.5331 | 1637 |
| 17 | KC405 | -93.4833 | 26.5706 | 1700 |
